# Population shifts in begomoviruses associated with tomato yellow leaf curl disease in western Mediterranean countries

**DOI:** 10.1101/2024.08.09.607290

**Authors:** Martine Granier, Mohamed Faize, Sandie Passera, Cica Urbino, Michel Peterschmitt

## Abstract

Tomato yellow leaf curl disease (TYLCD) was reported in western Mediterranean basin since the late 1980s. Based on intensive plant samplings performed in Spain, Italy and Morocco at different periods between the 1990s and 2014, several begomoviruses (family *Geminiviridae*) were identified as the cause of TYLCD. They comprise the native *Begomovirus solanumflavusardiniaense* (Tomato yellow leaf curl Sardinia virus, TYLCSAV), two strains of *Begomovirus coheni* (Tomato yellow leaf curl virus, TYLCV) introduced from the Middle East, and several types of TYLCV/TYLCSaV recombinants including the invasive recombinant TYLCV-IS76 in which the genome fragment inherited from TYLCSaV was unusually short. Although parental and recombinant TYLCD-associated begomoviruses were present in each country, country specificities were detected with respect to their relative prevalence, the infection profiles of individual tomato plants, and the recombination patterns of TYLCV/TYLCSaV recombinants. Considering geographic proximities and trade activities between these countries, and the efficient transmission of begomoviruses which are persistently transmitted by the polyphagous and tiny whitefly vector *Bemisia tabaci*, it was not known if these specificities would be maintained over time. To address this question, 105 tomato samples collected in the three countries between 2015 and 2019 were analysed with PCR tests previously designed to distinguish species, strains and recombinants of TYLCD associated begomoviruses, and with an original PCR test distinguishing emerging resistance-breaking recombinants bearing short TYLCSaV-inherited fragments like TYLCV-IS76 (Srec) from formerly reported recombinants with longer TYLCSaV fragments (Lrec). The results show that country specificities are still present, the most striking being the contrasted geographic distribution of Srec recombinants, with TYLCV-IS76 detected in Morocco but not in Italy, and TYLCV-IS141 and a new Srec recombinant (TYLCV-IMS60-2400) both detected in Italy and not in Morocco. Nevertheless, besides country specificities, a general population shift was revealed regarding TYLCV/TYLCSaV recombinants. Indeed, all the recombinant positive samples, irrespective of their geographic origin, were Srec-positive but Lrec-negative, which suggest that the emergence of Srec recombinants occurred at the expenses of the Lrec recombinants. These results are discussed in relation to the positive selection of Srec recombinants by *Ty-1* resistant tomato plants.

## 1. Introduction

Tomato (*Solanum lycopersicum* L.) is one of the New World (NW) originated vegetables that is presently grown worldwide including Mediterranean countries. During its global spread, it encountered Old World (OW) viruses and particularly viruses belonging to the genus *Begomovirus* of the family *Geminiviridae*. Indeed, according to the International Committee on Taxonomy of Viruses (https://ictv.global/taxonomy, ICTV_Master_Species_List_2023_MSL39), 112 Begomoviruses were isolated from tomato worldwide and most of them (69) from the OW. Noteworthy, a typical feature of OW begomoviruses is their ability to induce leaf curl, a feature which is highlighted in all the 69 names, as for example Tomato yellow leaf curl virus (TYLCV) or Tomato yellow leaf curl Sardinia virus (TYLCSaV). The names of the 43 NW begomoviruses indicate much more diverse symptoms like mosaic, mottle, distortion, rugose, streak… and only 8 of them were characterized by the terms “leaf curl” in their names.

Following the introduction of tomato in the western Mediterranean region, it encountered at least two OW begomoviruses, TYLCSaV and TYLCV. They both cause tomato yellow leaf curl disease (TYLCD), an economically important disease of tomato. TYLCSaV was reported first (Kheyr-Pour et al., 1991; Luisoni et al., 1989; Noris et al., 1994) and is generally considered to be indigenous to the Western Mediterranean region (Lefeuvre et al., 2010). TYLCV is a foreign begomovirus that probably arose somewhere in the Middle East (Lefeuvre et al., 2010). Two TYLCV strains were reported from western Mediterranean countries, TYLCV-Israel (TYLCV-IL) and the so-called mild strain, TYLCV-Mld. In Spain, the earliest plant samples in which TYLCV-Mld and TYLCV-IL were detected, were collected in 1997 (NavasCastillo et al., 1997) and 2002 (Morilla et al., 2003), respectively. In Italy, the earliest collected samples in which TYLCV was detected are from 2002 for the IL strain (Accotto et al., 2003) and 2012 for the Mld strain (Belabess et al., 2015). In Morocco, the earliest plant samples in which TYLCV-IL and TYLCV-Mld were detected, were collected in 1998 (Belabess et al., 2015; Peterschmitt et al., 1999). Although all three countries host TYLCD-associated viruses of the same species, country specificities were detected. For example, the displacement of TYLCSaV induced by the arrival of TYLCV was different between Spain and Italy. Indeed, whereas in Spain the displacement was virtually total, in Italy, virus populations reached an equilibrium in which the two virus species coexist (Davino et al., 2006; Davino et al., 2009; Garcia-Andres et al., 2007; SanchezCampos et al., 1999).

Since the spread of TYLCV into western Mediterranean countries, the diversity of begomoviruses has increased considerably, and not only because of the subsequent introduction of a non-native virus species from Asia, i.e. *Tomato leaf curl New Delhi virus* (ToLCNDV) (Juárez et al., 2013), but also due to recombinants between TYLCV and TYLCSaV as reported from Spain (Garcia-Andres et al., 2006; Monci et al., 2002), Italy (Belabess et al., 2015; Davino et al., 2008; Granier et al., 2019; Panno et al., 2018) and Morocco (Belabess et al., 2015).

The earliest collection of plant samples detected positive for TYLCV/TYLCSaV recombinants was in 1998 in Spain and Morocco (Belabess et al., 2015; Monci et al., 2002) and 2002 in Italy (Davino et al., 2008). Although the collection dates of these recombinant-positive samples were fairly similar, the earliest dates at which plants were detected with only recombinants, i.e. testing negative for parental viruses, are very different. Indeed, whereas in Spain such plants were detected 1 year after the first detection of recombinants (Monci et al., 2002), in Morocco and Italy the delay was at least 10 years (Belabess et al., 2015). In natural conditions, when a recombinant is detected in a plant without parental viruses, it generally means that it is fit enough to spread autonomously from plant to plant and is potentially emerging. It seems therefore that the agroecological environment of Spain was more conducive to the emergence and spread of TYLCV/TYLCSaV recombinants than the Italian and Moroccan environment. The contrasted practice of bean (*Phaseolus vulgaris* L.) cultivation between Southern Spain and Sicily was suggested to be critical (Davino et al., 2009); in Southern Spain, this host of TYLCD-associated viruses is regularly cultivated between tomato crops, but not in Sicily. Thus, this specific cultivation practice of beans may have favoured the emergence of the bean adapted recombinant Tomato yellow leaf curl Malaga virus (TYLCMalV) in Spain (Monci et al., 2002).

Another country specificity is related to the recombination profiles of TYLCV/TYLCSaV recombinants detected in plants tested negative for parental-type viruses reflecting their autonomous spread. The two autonomously spreading recombinants reported from Spain, i.e., TYLCMalV (Monci et al., 2002) and Tomato yellow leaf curl Axarquia virus (TYLCAxV) (Garcia-Andres et al., 2006) have both inherited a long TYLCSaV fragment of at least 1000 nts, including at least the two viral strand-encoded genes V1 and V2 and their promoter region up to the origin of replication. Although this type of recombinants were also reported from Italy (Davino et al., 2012) and are suspected from Morocco (Belabess et al., 2015), they were not found to spread autonomously. After 2000, TYLCD-resistant-tomato cultivars bearing the *Ty-1* resistance gene have progressively replaced susceptible cultivars. This replacement coincided with the emergence of the autonomously spreading recombinant TYLCV-IS76 in Morocco that displaced its parental viruses (Belabess et al., 2015). This coincidence together with a relatively higher experimental competitiveness of TYLCV-IS76 in *Ty-1* resistant plants (Belabess et al., 2016) is consistent with a *Ty-1*-resistance driven emergence of TYLCV-IS76 by its positive selection. Intriguingly, although *Ty-1* cultivars were similarly deployed in Spain and Italy, TYLCV-IS76 did not emerge in these countries; TYLCV-IS76 recombinants were reported in Spain (Fortes et al., 2023; Torre et al., 2019), but due to their high nucleotide identity with Moroccan representatives, they presumably originate from Morocco (Torre et al., 2018). However, in Italy, a similar recombinant emerged, TYLCV-IS141. Its detection from *Ty-1*-resistant plants (Granier et al., 2019; Panno et al., 2018) and its relatively higher experimental competitiveness in *Ty-1* plants (Urbino et al., 2020) suggest that its emergence was triggered by the deployment of *Ty-1* resistant cultivars. In comparison to the autonomously spreading recombinants that emerged in Spain, those that emerged in Italy and Morocco inherited a much shorter TYLCSaV fragment, located between the origin of replication and the 5’ end of the V2 gene. Intriguingly, while the emerging recombinants reported from Italy have a TYLCSaV-inherited fragment of 141 nts (Belabess et al., 2015; Granier et al., 2019; Panno et al., 2018), the emerging recombinant reported from Morocco has only a 76 nts fragment inherited from TYLCSaV (Belabess et al., 2015). These recombinants were named TYLCV-IS76 and TYLCV-IS141 according to the position of their typical recombination breakpoints (Belabess et al., 2015; Belabess et al., 2018)..

Considering geographic proximity and trade activities between Morocco, Italy and Spain, and the efficient and persistent transmission of these viruses with the tiny polyphagous winged whitefly vector *Bemisia tabaci*, it was not known if these specificities would be maintained or rather fade over time due to natural or anthropogenic-induced virus spread. To address this question, we analysed the infection profiles of tomato samples collected from the three countries between 2015 and 2019. The infection profile of each plant was first determined with the multiplex PCR tests described by Belabess et al (Belabess et al., 2015). Then, to monitor more specifically the prevalence of recombinants with a resistance breaking potential like TYLCV-IS76 and TYLCV-IS141, a new test was designed to distinguish such recombinants characterized by their short TYLCSaV inherited fragment (Srec), from those that inherited longer TYLCSaV fragments like TYLCMalV and TYLCAxV (Lrec). Although the results show that country specificities are maintained with contrasted prevalence of recombinant and parental type viruses between countries, they also show a general population shift with the emergence of Srec recombinants at the expenses of the Lrec recombinants not detected in any of the samples from the three countries. These results are discussed in relation to the deployment of Ty-1-resistant cultivars and their positive selection of Srec recombinants.

## 2. Material and Methods

### 2.1. Plant material

Leaf samples were collected from net-house protected tomato crops between 2015 and 2019 in Morocco, Spain and Italy (Table 2, Supplementary material 1).

In Morocco, 3 samples were collected in December 2015 in Agadir, and 48 samples were collected in 2017 from 6 locations along the Atlantic coast between Casablanca and Agadir. These 48 samples were from 12 farms, with 4 samples per farm collected on tomato plants exhibiting TYLCD symptoms. Collection sites from North to South were Azmour with 1 farm sampled in April, Chtouka ouled laissaoui with one farm sampled in March, Ouled Frej with 3 farms sampled in February, Oualidia with 4 farms sampled in March, Safi doukkala had lhrara with one farm sampled in March, and Agadir with 2 farms sampled in January.

In Italy, 37 samples were collected between 2015 and 2019. Four samples are from Sardinia and 33 samples from Sicily. Samples from Sardinia were previously reported (Granier et al. 2019) but were further analysed here with a PCR test distinguishing Lrec and Srec type recombinants.

In Spain, a total of 17 samples were collected between 2015 and 2018 in the South-East of the country within the tomato-growing regions of Murcia and Almeria

### 2.2. DNA extraction

The 48 leaf samples collected in 2017 in Morocco were lyophilized and total DNA was extracted with a reported protocol (Dellaporta et al., 1983) modified as follows. The amount of lyophilized plant material used per extraction ranged between 1.25 and 20 mg. Plant material was ground in 400 μL extraction buffer (100 mM Tris-HCl pH 8.0, 50 mM EDTA, 500 mM NaCl, 1% SDS, 36 mM Na2SO3, and 0.1 mg/ml RNase), incubated at 65°C for 10 min and centrifuged (16,000 g, 10 min). One volume of isopropanol was added to 300 μL of the supernatant and nucleic acids were precipitated by centrifugation (16,000 g, 20 min); the pellet was washed with ethanol 70% and then resuspended in 50 μL water. As PCR amplifications failed for 32 of the samples, total DNA was extracted again for these 32 samples using a silica-based DNA extraction kit (DNeasy Plant Mini Kit, QIAGEN S.A., Courtaboeuf, France) with 10 mg of lyophilized plant material per extraction.

Total DNA from all the other samples from Morocco, and those from Italy and Spain was extracted with the silica-based DNA extraction kit “NucleoSpin Plant II, Mini kit for DNA from plants” (Macherey Nagel SAS, Hoerdt, France), except for the samples from Sardinia for which the Qiagen-DNeasy Plant Mini kit was used.

### 2.3. PCR tests and primer design

Plant samples were first analysed with the two multiplex PCR tests MxPCR1 and MxPCR2, with which the presence of TYLCV-IL, TYLCV-Mld, TYLCSaV, and recombinants derived from them (Belabess et al., 2015) can be specifically detected. Due to the fact that all reported TYLCV/TYLCSaV recombinants bear a recombination breakpoint in the stem loop bearing the origin of replication (OR), the test is expected to detect all recombinants between TYLCV-IL and TYLCSaV and between TYLCV-Mld and TYLCSaV. Another typical feature of these reported recombinants is that the sequence contiguous to OR and located at the complementary sense gene side IS always inherited from TYLCV, whereas the sequence located at the other side is always inherited from TYLCSaV; reciprocal recombinants have not been reported so far.

Samples detected positive for the presence of TYLCV/TYLCSaV recombinants with MxPCR1, were analysed further with original PCR tests designed in this study to distinguish between recombinants that inherited a short TYLCSaV fragment (Srec) like TYLCV-IS76 or TYLCV-IS141, from those that inherited long TYLCSaV fragments (Lrec) like TYLCAxV (Garcia-Andres et al., 2006) or TYLCMalV (Monci et al., 2002).

The pair of PCR primers targeting Srec recombinants, comprise the previously reported forward primer Sar19F (Belabess et al., 2018), complementary to TYLCSaV genome sequences, and an original reverse primer IM650R, complementary to TYLCV genome sequences (Fig 1A and Table 1). The previously designed reverse primers Ty153R and Ty282R (Belabess et al., 2018) were used to further analyse Srec-positive samples for the presence of IS76- or IS141-type recombinants. The original forward PCR primer targeting Lrec recombinants is Sar599F, a primer complementary to TYLCSaV genome sequences (Fig 1C and Table 1). To distinguish Lrec having a TYLCV-IL parental virus like TYLCAxV (Garcia-Andres et al., 2006) from Lrec having a TYLCV-Mld parental virus like TYLCMalV (Monci et al., 2002), two original reverse primers were used, IL-2258R and Mld-2310R, respectively (Table 1). Noteworthy, as the TYLCV-IL and TYLCV-Mld genomes can be distinguished only between position 2000 and the end of the genome, it was not possible to infer from the Srec PCR products, located between position 1 and 650, whether the TYLCV parent of the Srec was of the IL or the Mld strain. The specificity of the five original primers of the Srec/Lrec PCR tests (Sar599F, IM650R, IL2258R and Mld2310R), was determined in silico with full genomic sequences available in GenBank, 121 from TYLCV-IL clones, 18 from TYLCV-Mld clones and 21 from TYLCSaV clones. The nucleotides determining the specificity of the primers were written with bold letters in Table 1. Noteworthy, some of the primers were degenerated to take into account the intra-species nucleotide diversity.

**Figure 1:**
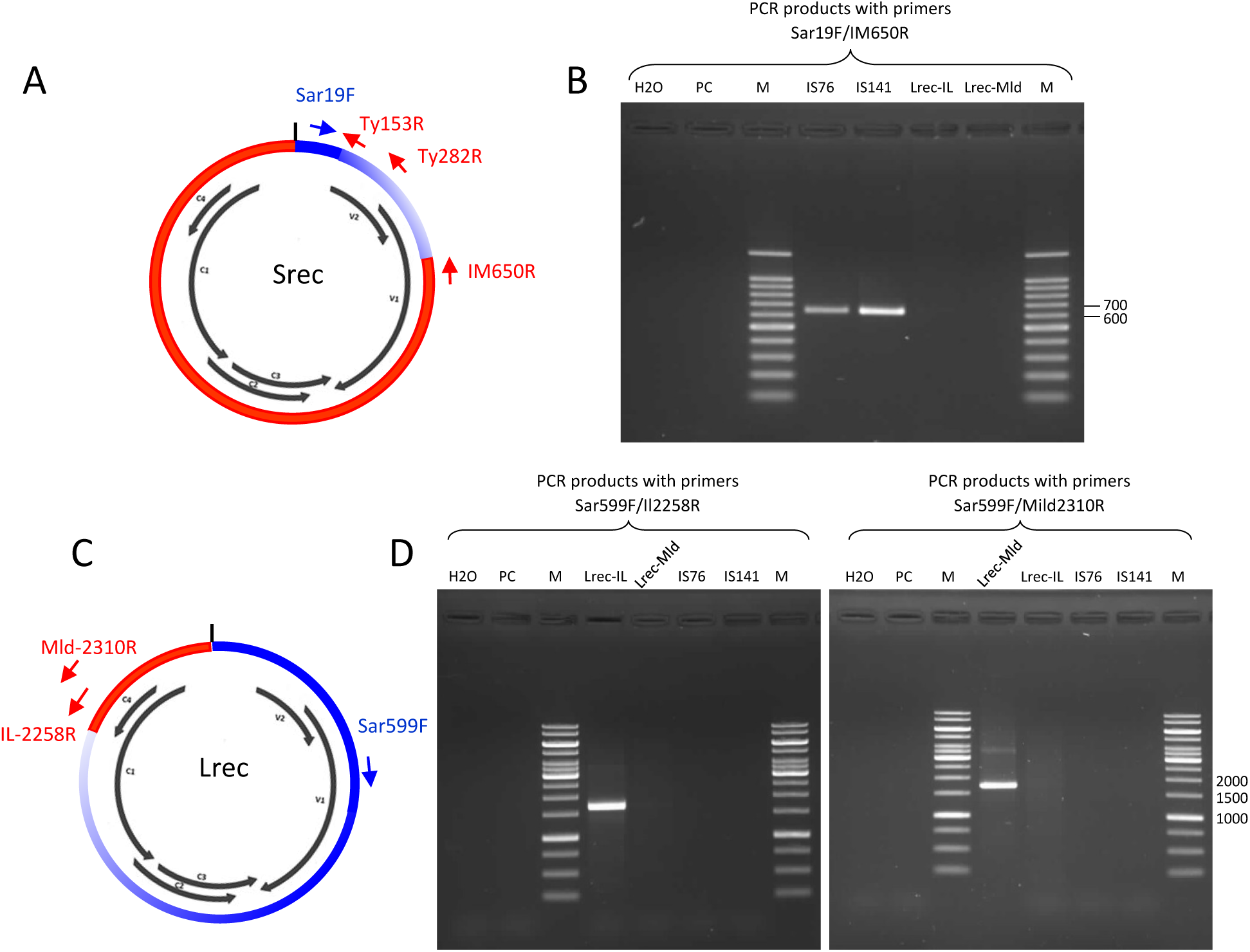
Presentation of PCR primers designed to distinguish Srec and Lrec recombinants. (A) PCR primers targeting recombinants that inherited short TYLCSaV derived fragments of less than 623 nts (Srec). Primer positions are schematically presented on a circular viral genome; black arrows correspond to the commonly reported open reading frames and the short vertical line at the top of the circle correspond to the stem loop bearing the origin of replication. (B) PCR products obtained with Srec-targeting primers resolved on a 1.5% agarose and stained with ethidium bromide (M, Benchtop ladder, Promega, France). (C) PCR primers targeting recombinants that inherited TYLCSaV-derived fragments of more than 623nts (Lrec). The reverse primer IL-2258R, specifically hybridize to Lrec having a TYLCV-IL parental virus (Lrec-IL), whereas Mld-2310R specifically hybridize to Lrec having a TYLCV-Mld parent (Lrec-Mld). (D) PCR products obtained with Lrec-targeting primers resolved on a 1% agarose gel (M, Gene ruler 1kb, Fisher Scientific SAS, France). All PCR primers are described in Table 1. IS76, TYLCV-IS76; IS141, TYLCV-IS141; PC, total DNA from a mock inoculated plant.

**Table 1:**
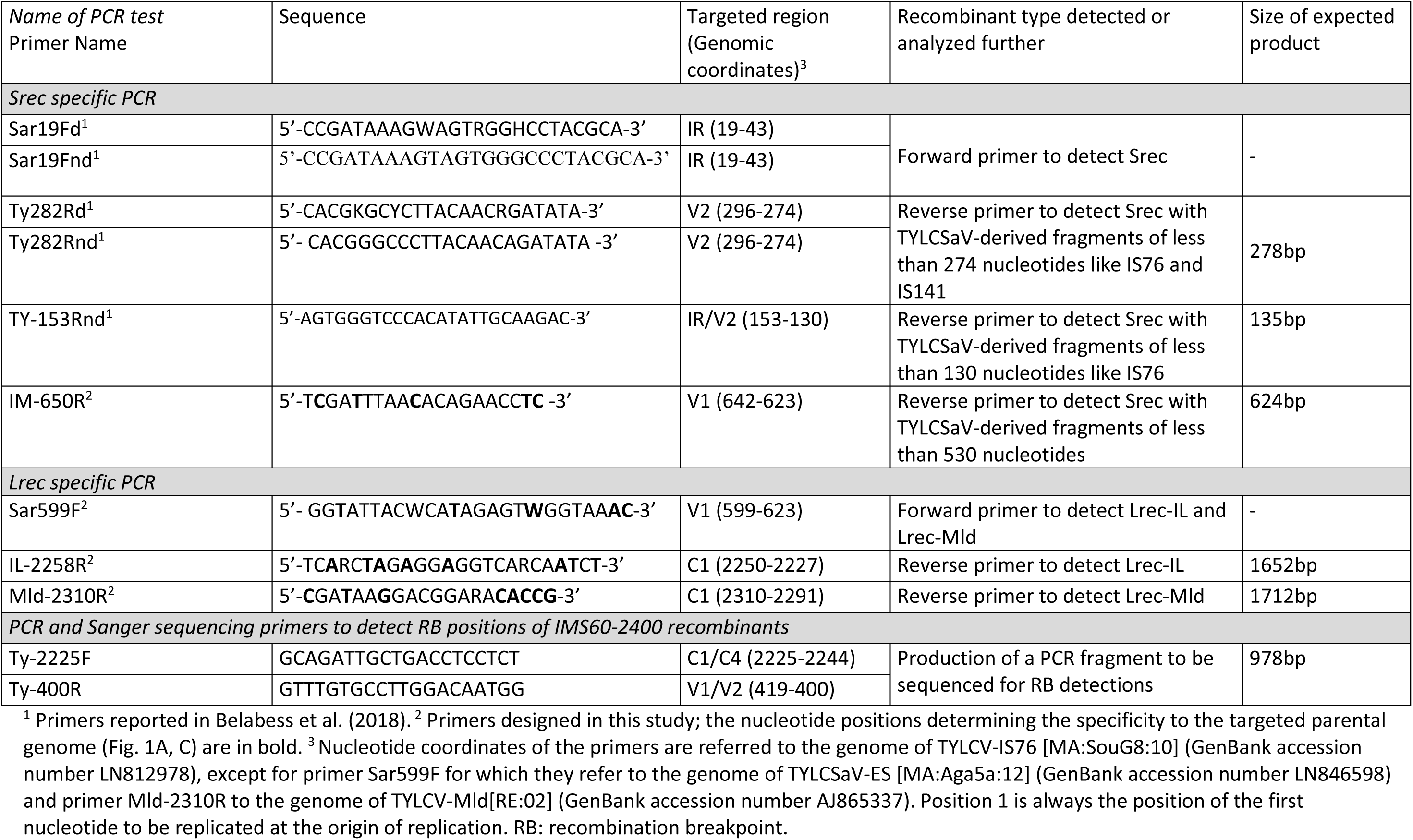
Description of PCR primers and primer products.

Considering the position of the primers described above, the Srec recombinants are TYLCV/TYLCSaV recombinants that inherited a TYLCSaV fragment starting from the OR and extending at the maximum up to position 622, just before the 3’ terminal nucleotide of the TYLCV specific primer IM650R (Table 1). Lrec recombinants are TYLCV/TYLCSaV recombinants that inherited a TYLCSaV fragment that extended at least up to position 623, corresponding to the 3’ terminal nucleotide of primer Sar599F. Lrec-IL are Lrec recombinants for which the TYLCV parent is from the IL strain whereas Lrec-Mld are Lrec recombinants with a TYLCV-Mld parent.

The cycling protocol for the amplification of Srec recombinants was as follows: 5 min denaturation at 95 °C, followed by 30 cycles, each consisting of 45 s at 95 °C, 45 s at −52 °C with reverse primers IM650R and TY282Rd, 54°C with primer TY282Rnd and 55°C with primer TY153Rnd - and 45 s elongation at 72°C. The PCR program was ended by a 5- or 10-min incubation at 72 °C. The PCR products were resolved by electrophoresis in a 1% to 2% agarose gel according to the different lengths of the expected fragments, and visualized with ethidium bromide staining.

The cycling protocol for the amplification of Lrec recombinants was as follow: 5 min denaturation at 95 °C, followed by 30 cycles, each consisting of 45 s at 95 °C, 45 s at 50 °C, and 90 s at 72 °C. The PCR program was ended by a 10 min incubation at 72 °C. The same protocol was used for Lrec-IL and Lrec-Mld. The PCR products were resolved by electrophoresis in a 1% agarose gel and visualized with ethidium bromide staining.

The Srec/Lrec discriminating PCR tests were validated with total DNA extracted from tomato plants agroinoculated with infectious viral clones. Two DNA extracts were used as positive controls for Srec, one from a tomato plant of the cultivar Pristyla (Gautier Semences, France) agroinfected with TYLCV-IS76 (Belabess et al., 2015), and one from the cultivar Calvi (Gautier Semences) agroinfected with TYLCV-IS141 (TYLCV-IS141[IT:Sic1; 13]) (Urbino et al., 2020). The positive control for Lrec-IL was an extract of total DNA from a tomato plant of the cultivar Pristyla co-agroinoculated with TYLCV-IL and TYLCSaV (Belabess et al., 2018); this DNA was previously extracted from a leaf sample collected from plant R11, 180 days post inoculation, and the presence of Lrec recombinants in this sample was determined by Sanger sequencing of a PCR product generated with primers designed to detect TYLCV/TYLCSaV recombinants [see Fig. 8 and Material and Methods of (Belabess et al., 2018)]. The positive control for Lrec-Mld was a recombinant plasmid DNA containing the full-length genome of a TYLCVMalV clone (Genbank accession number LN846611) (Belabess et al., 2015); the viral clone was obtained from a bean plant (H5) sampled in March 2012 in the Souss region of Morocco. As negative plant control, we used total DNA extracted from a tomato plant of the cultivar Pristyla infiltrated with an agrobacterium clone carrying an empty PC2300 binary vector. The specificity of each primer pair was tested with DNA extracts of target and non-target recombinants, and with the negative plant control.

### 2.4. Sequencing of PCR products and sequence analysis

Some of the samples in which Srec type recombinants were detected by PCR, were further analysed by sequencing to determine non-OR recombination breakpoints. To do this, the PCR products generated with the Srec-specific PCR primer pair Sar19/IM-650R or the products generated with the primer pair Ty-2225F/Ty-400R were sequenced by Sanger sequencing (Eurofins Genomics, Ebersberg, Germany) using the PCR primer IM650R, and both PCR primers in the case of the second PCR product (Table 1).

Additional sequencing of PCR products was done for plant samples detected positive for TYLCV but only for those collected in Spain. Two TYLCV genotypes, group 1 and group 2, were previously identified from tomato plants sampled in Southeast Spain in 2015 and 2016 (Torre et al., 2018). Group 1 viruses are related to the two early TYLCV-IL isolates of Southeast Spain, both sampled on pepper (*Capsicum annuum*), one in 1999 in Almeria (TYLCV-IL[ES:Alm:Pep:99], GenBank accession number AJ489258) (Morilla et al., 2005) and one in 2003 in Malaga (TYLCV-IL [ES:Mlg:TY11:Pep2003], GenBank accession number KC953602). Group 2 viruses are closely related to two TYLCV-IL isolates sampled on tomato in Morocco, TYLCV-IL[MA:El Jadida:Tom:2013] (GenBank accession number LN846613 and LN846613) (Belabess et al., 2015). The most parsimonious evolutionary scenario suggested that the TYLCV isolates of Group 2 are back recombinant isolates derived from TYLCV-IS76 (Torre et al., 2018). The relatedness to group 1 or 2 of the TYLCV isolates from Spain, was based on a 803-nucleotide fragment amplified with the forward primer IL-2629F (5’-GGTGTCCCTCAAAGCTCTAWG-3’) of the multiplex PCR test MxPCR2 (Belabess et al., 2015) and the original reverse primer IM650R (Table 1). Using IM-650R as sequencing primer, 11 group 1/group2 discriminating nucleotide positions could be determined. Sequences were aligned with 27 homologous genomic sequences from TYLCVs for which complete viral genomes are available in Genbank, 25 from Spain and the 2 from Morocco representative of group 2. The alignment was performed with the Optimal Alignment method of DNAMAN (version 5.0; Lynnon BioSoft, Quebec, Canada). A phylogenetic tree was set up with a Jukes and Cantor distance matrix using the neighbour-joining method (Saitou and Nei, 1987) as implemented in the DNAMAN software. The sequences used for the alignment are described on the phylogenetic tree presented in the section Results.

### 2.5. Cloning and sequencing of complete viral genomes

A new Srec type recombinant bearing a recombination breakpoint at position 60 was detected in Sicily by sequencing PCR products. To characterize the full-length genome of this new type of Srec recombinant, it was cloned from total DNA extracted from three plant samples with the DNeasy Plant Minikit. Circular DNA molecules were amplified using a TempliPhi Amplification kit (GE Healthcare, UK) according to the manufacturer’s instructions, and digested with restriction enzymes that are expected to be potentially unique according to reported sequences of parental type genomes of TYLCV and TYLCSaV. Digestion of RCA products with *Sac*I generated linear fragment of about 2.8kb, a size expected with full-length begomovirus genomes. The restricted fragments were ligated into the plasmid pBC and ligation products were introduced into the bacterial strain *Escherichia coli* DH5alpha. The cloned fragments were sequenced with M13 primers and virus-specific primers (not communicated). Sequences were analysed with Chromas (Technelysium Pty Ltd, Australia) and DNAMAN.

### 2.6. Recombination analysis

A recombination analysis was performed on full-length genomes of the new Sicilian Srec recombinant bearing a recombination breakpoint at position 60. Recombination analysis was performed with the Recombination Detection Program RDP4.101 (Martin et al., 2015) with default settings. The genomes used for the input alignment are from clones of potential parental viruses present in Sicily, i.e., TYLCV-IL, TYLCV-Mld, TYLCSaV and the recombinant TYLCV-IS141. The genomes of these viruses were from clones isolated either from Italy or nearby countries in the Western Mediterranean region. The genomes used for the alignment are from 17 TYLCV-IL clones (GenBank accession numbers AJ489258, DQ144621, EF060196, KC953602, LN846613, LN846615, LN846616, LN846617, MH644779, MH644782, MH644786, MH644787, MH644788, MH680947, MH680949, MH680953, MH680958), 3 TYLCV-Mld clones (AF071228, AF105975, AJ519441), 16 TYLCSaV clones (AJ519675, AY702650, AY736854, GU951759, JN859134, JN859136, KC 953603, KC953604, LN846595, LN846596, LN846597, LN846598, L27708, X61153, Z28390, Z25751) and 3 TYLCV-IS141 clones (MH817479, MG489968, MF405078); The genome of a TYLCV-Mld clone (GenBank accession number KJ913683) isolated in Dominican Republic was added to the alignment because it showed the highest genetic relatedness with the RB60-bearing recombinant according to preliminary analysis. A similarity plot was produced with the application SimPlot++ (Samson et al., 2022) to show the complete recombination profile of the RB60 bearing recombinant.

## 3. Results

### 3.1. Srec and Lrec recombinants can be distinguished with a discriminating PCR test

An original PCR test was designed to distinguish TYLCV/TYLCSaV recombinants that inherited short TYLCSaV derived fragments of less than 623 nts (Srec), from those that inherited TYLCSaV-derived fragments of more than 623nts (Lrec).

Amplification products generated with Srec/Lrec discriminating PCR primers are presented in Fig. 1B and Fig. 1D. Srec primers do not generate any amplification product with Lrec infected plants and vice versa. No amplification product was detected with a mock-inoculated plant infiltrated with the agrobacterium clone carrying an empty PC2300 binary vector. A PCR product of higher-than-expected size was resolved for the positive control of Lrec-Mld (Fig. 1D). As the control DNA for Lrec-Mld is a full-length viral genome cloned in an *Escherichia coli* plasmid, we suppose that this long product was generated with elongations that went all around the recombinant plasmid.

The Srec/Lrec discriminating PCR test was subsequently used to analyse tomato samples detected positive for the presence of TYLCV/TYLCSaV recombinants according to the results obtained with the MxPCR1 test (Belabess et al., 2015) targeting recombination breakpoints at the origin of replication.

### 3.2. Infection profiles of samples collected in Morocco

Among the 51 samples analysed, 46 were detected positive for at least one PCR-targeted virus and three infection profiles were distinguished (Table 2): TYLCSaV (1 sample), IL/Sar (43 samples), and coinfection with IL/Sar and Mld/Sar (2 samples). Moroccan samples can be characterized as follows. (i) The parental virus types TYLCV-IL and TYLCV-Mld were not detected and the TYLCSaV type was detected in only one sample. (ii) All the 46 PCR-positive samples, were positive for IL/Sar except the sample detected positive for TYLCSaV only. (iii) The recombinant-positive samples were negative for recombinant-types that inherited long TYLCSaV fragments (Lrec), but positive for Srec recombinants. (iv) The recombination breakpoints within the sequence of the Srec PCR products were analysed for 5 plant samples by sequencing. For representativity reasons, the sequencing was performed on samples of different regions and collection years. Recombination breakpoints were shown to be all at position 76, like that of the typical recombination breakpoint of TYLCV-IS76.

**Table 2:**
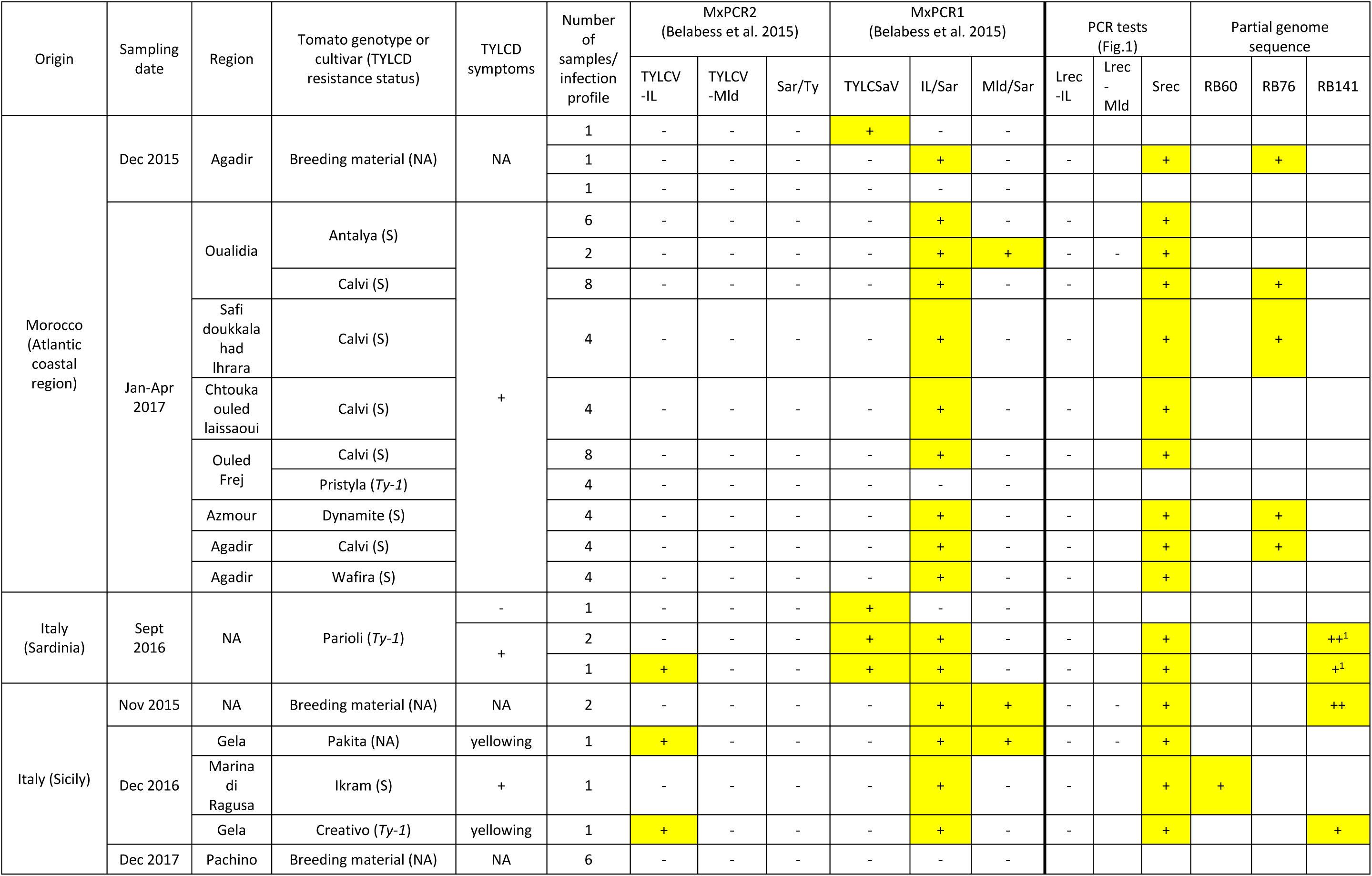
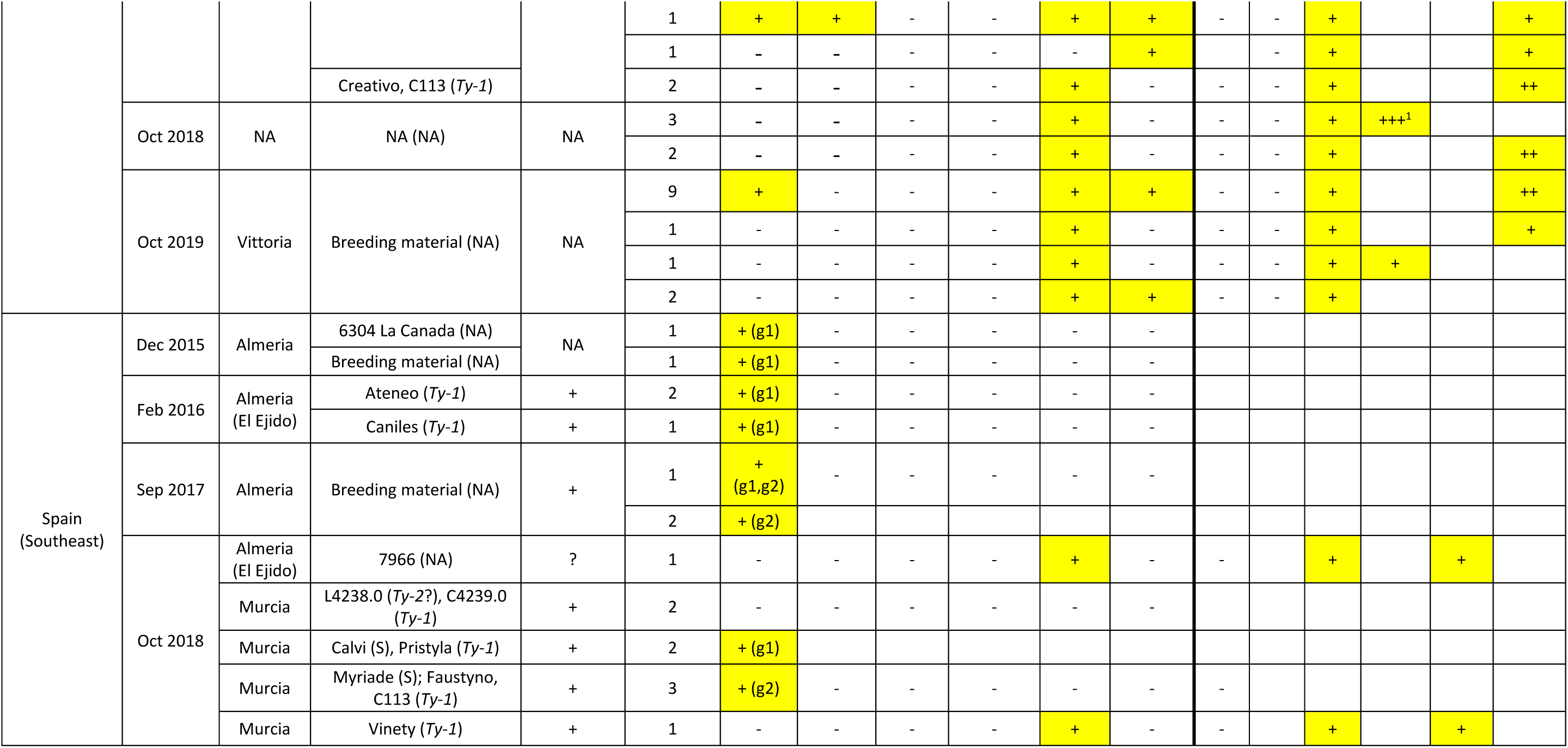
Begomovirus infection profiles of tomato plants sampled in Morocco, Italy and Spain. Total DNA of plant samples were analysed by PCR and DNA sequencing. The presence of Tomato yellow leaf curl virus (TYLCV), Tomato yellow leaf curl Sardinia virus (TYLCSaV) and the interspecies recombinants derived from them were tested with the multiplex PCR MxPCR1 and MxPCR2 (Belabess et al., 2015); note that the test distinguishes the Israel and Mild strain of TYLCV and the recombinants derived from them, TYLCV-IL/TYLCSaV (IL/Sar) and TYLCV-Mld/TYLCSaV (Mld/Sar). IL/Sar and Mld/Sar correspond to recombinants in which the sequence contiguous to the origin of replication on the virus sense gene side is inherited from TYLCSaV. Inversely, “Sar/Ty” corresponds to potential reciprocal recombinants TYLCSaV/TYLCV. Samples from Spain detected positive for TYLCV-IL were analysed further by sequencing to specify to which formerly identified group they may belong to (Torre et al., 2018), i.e., group 1 (g1) or group 2 (g2) as shown in Fig.4. Samples detected positive for recombinants were analysed further by PCR tests to distinguish them according to the size of the fragment inherited from TYLCSaV, either long (Lrec-IL or Lrec-Mld) or short (Srec) (Fig. 1, Table 1). Positions of recombination breakpoints were identified by sequencing PCR products (last 3 columns); the number of “+” indicates the number of samples that were analysed by sequencing. Recombination breakpoints (RB) were detected at position 60, 76, or 141. In the columns corresponding to PCR tests, + and - indicate that a PCR product of the expected size was detected or not detected, respectively; an empty box means that the PCR test was not done. The boxes containing a “+” were coloured in yellow to facilitate the reading of the table. In the column describing the samples, NA means not available. The TYLCD resistance status of tomato genotypes or cultivar is described with the presence of the resistance genes *Ty-1* and *Ty-2*, and with S meaning susceptible. ^1^Samples for which recombinant genomes were totally sequenced following cloning; the three sequences of the Sardinian TYLCV-IS141 were reported previously (Granier et al., 2019).

The 4 samples collected in Ouled-Frej from symptomatic *Ty-1* resistant tomato plants of the cultivar Pristyla tested negative for all targeted begomoviruses whereas all the 8 samples from the susceptible cultivar Calvi were detected positive. It is possible that the PCR tests were not sensitive enough to detect the low virus amounts of Ty1 resistant Pristyla plants which were reported to be about 10 times lower than the amounts in nearly isogenic susceptible tomato plants (Belabess et al., 2016). This hypothesis is consistent with the observation of a relatively faint band observed in lane IS76 of the agarose gel shown in Fig. 1A, corresponding to an amplification product obtained with the Srec specific primers, from a Pristyla tomato plant agroinoculated with a clone of TYLCV-IS76. It is also possible that the milder TYLCD symptoms exhibited by *Ty-1* resistant plants may have been confused with other types of symptoms. Finally, it cannot be excluded that these samples are infected with TYLCD-associated begomoviruses that are not detectable with the PCR approach used.

### 3.3. Infection profiles of samples collected in Italy

Among the 37 samples analysed, 31 were detected positive for at least one PCR-targeted virus. The infection profiles are relatively heterogeneous. Indeed, unlike in Morocco where only one parental-type virus (TYLCSaV) and only one type of TYLCV/TYLCSaV recombinant (TYLCV-IS76) were detected, in Italy, all three parental type viruses were detected as well as two types of recombinants, TYLCV-IS141 and a new one described below, TYLCV-IMS60-2400 (IMS60-2400). Moreover, whereas only three infection profiles were detected in Moroccan samples, up to 9 infection profiles were detected in Italian samples. In spite of these differences between infection profiles of Italian and Moroccan samples, some common features can be noted. Thus, like in Morocco, only one sample was infected with a parental virus alone (TYLCSaV), whereas all the other virus-positive samples, were positive for the presence of IL/Sar or Mld/Sar recombinants. Moreover, like in Morocco, all the recombinant-positive samples are negative for Lrec recombinants, but positive for Srec recombinants.

To further characterize the Srec recombinants, the Srec PCR products from 20 of the 30 Srec-positive samples were analysed by sequencing; for representativity reasons, samples were selected from each collection year and from each reported sampling locations. The sequencing results revealed that 15 samples contained Srec recombinants of the IS141-type with a recombination breakpoint around position 141 (RB141), whereas 5 samples contained a new type of Srec recombinants with a recombination breakpoint located within the 57 to 62 position-range of identical nucleotides between TYLCV and TYLCSaV parental genomes (Fig. 2A). The recombination breakpoint was named RB60, 60 being one of the two median positions of the 6 non-discriminating nucleotide range in which recombination events occurred. Three RB60-bearing recombinant genomes were cloned from three samples collected in October 2018 and were fully sequenced (GenBank accession numbers OR473055, OR473056, OR473057). The genomes are 2783 or 2784 nts in length and differ from each other at 1, 8 or 9 positions. The potential presence of other recombination breakpoints was analysed with the program RDP4. Two recombinant regions were detected (Table 3). One was detected in the intergenic region of the viral sense gene side between RB60 and the stem loop bearing the origin of replication known as a recombination hot spot in begomoviruses (Lefeuvre et al., 2007). A longer one was detected between the stem loop and an RB at position 2400 (RB2400) including the 5’ terminal regions of the C1 and C4 genes and the intergenic region on the complementary sense gene side. Considering the low probabilities of false positives obtained with respectively 5 and 7 methods of recombination detection in RDP4, the recombinant origin of these two regions was strongly supported. The new recombinant was named IMS60-2400 in which “I” stand for the IL strain of TYLCV, the major parent, and “M” and “S” for the minor parents, i.e., the Mild strain of TYLCV and TYLCSaV; 60 and 2400 refer to median nucleotide positions within the non-discriminating regions in which the two non-OR recombination events occurred (Fig. 2A, 2B). The two samples detected positive for RB60-bearing recombinants by partial sequencing but for which full-length genomes were not cloned and sequenced (Table 2), were shown to contain both the RB60 and RB2400 by sequencing a PCR product generated and sequenced with the primer pair Ty-2225F/Ty-400R (Table 1).

**Figure 2:**
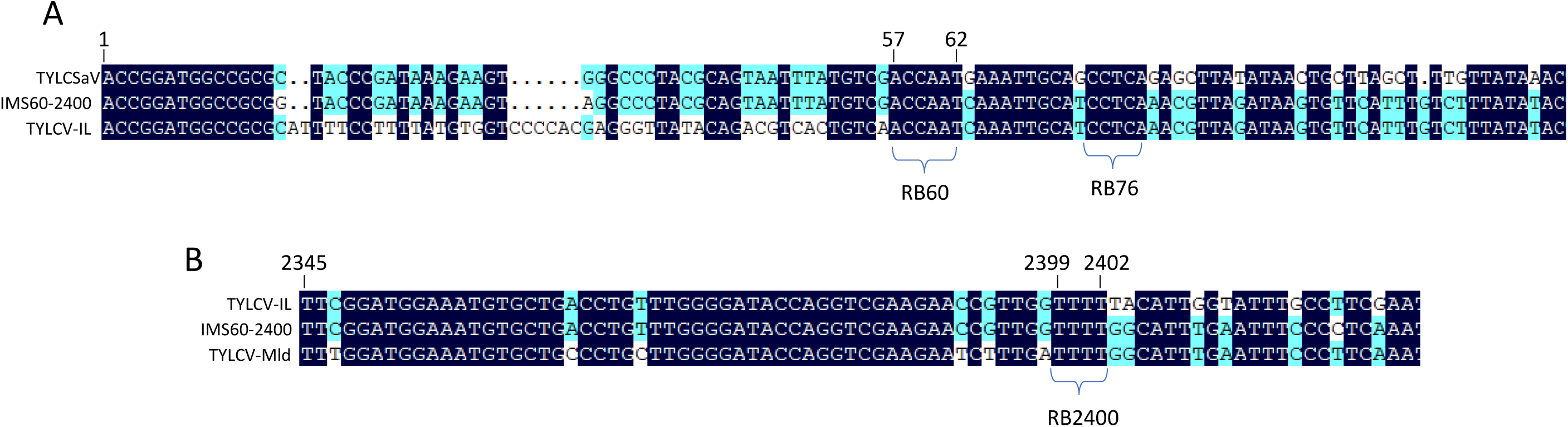
DNA alignments showing the typical recombination breakpoints of the TYLCV-IMS60-2400 recombinant. (A) Alignment showing the recombination breakpoint RB60. The partial DNA sequences are from the genome of three viral clones, TYLCV-IMS60-2400 (GenBank accession number OR473055) TYLCSaV-Sic from Tunisia (GenBank accession number AY736854), and TYLCV-IL from Italy (GenBank accession number DQ144621). Number 1 is the nucleotide position of the origin of replication, and positions 57 to 62 is the range of non-discriminating nucleotides bearing the recombination breakpoint RB60. The range of non-discriminating positions bearing RB76 is shown for information. (B) Alignment showing the recombination breakpoint RB2400. The partial DNA sequences are from the genome of three viral clones, TYLCV-IMS60-2400, TYLCV-IL and TYLCV-Mld from Dominican Republic (GenBank accession number KJ913683). Positions 2399 to 2402 is the range of non-discriminating nucleotides bearing the recombination breakpoint RB2400. Nucleotide positions are numbered according to the genome of TYLCV-IMS60-2400.

**Figure 3:**
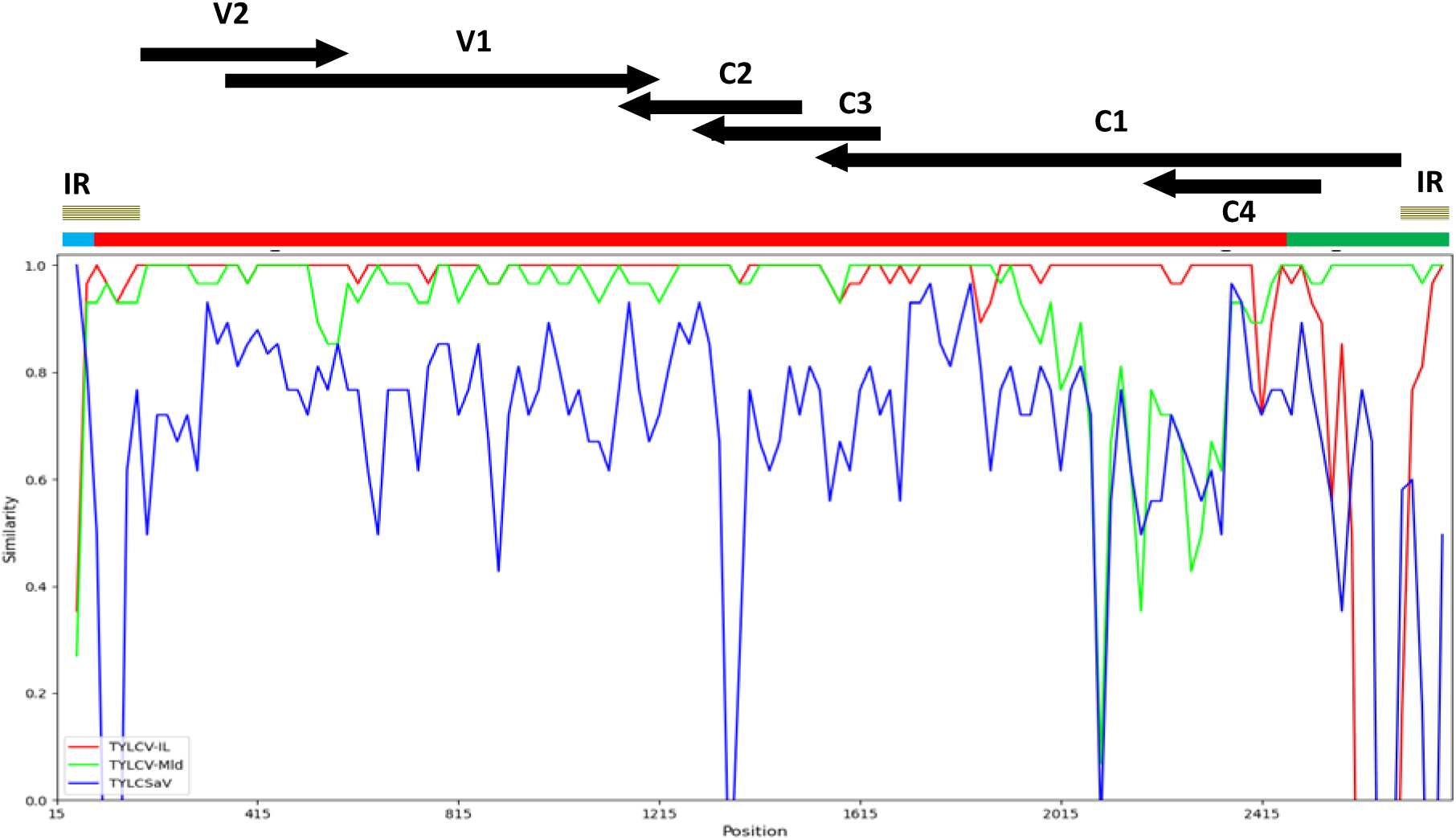
Similarity plot showing the recombination profile of TYLCV-IMS60-2400. The similarity plot was produced with the SimPlot++ application. The distance between sequences was determined with the Jukes Cantor model within a 30-nucleotide window slid by 20-nucleotide steps along the genome. The parental viruses were those inferred from the RDP analysis (Table 3). The recombination profile of the IMS60-2400 recombinant is presented in a linear form with the horizontal bar at the top of the similarity plot. The parental origin of the recombinant regions is presented with the colour code used in the similarity plot. The intergenic region and the six commonly reported ORFs of monopartite begomoviruses are shown at the top of the figure.

**Table 3:** Detection of recombinant regions in the genome of TYLCV-IMS60-2400. (Genbank accession number OR473055) with methods available in RDP4: RDP (R), GENECONV (G), BOOTSCAN (B), MAXIMUM CHI SQUARE (M), CHIMAERA (C), SISTER SCAN (S) and 3SEQ (3S). The reported P-value is for the method in bold type and is the best P-value of the detected region. Upper-case letters were used for methods detecting the potential recombinant region with a false positive probability of less than 10^-9^, whereas lower-case letters were used when probability is between 10^-4^ and 10^-2^. The inferred recombinant regions are characterized by the positions of their terminal nucleotides; positions are according to the genome of TYLCV-IMS60-2400, position 1 being the origin of replication. Genes and intergenic regions contained in the recombinant regions are indicated. IR, intergenic region.

| Positions of the recombinant region | Genomic regions | Major parent | Minor parent | Detection methods | P value |
| --- | --- | --- | --- | --- | --- |
| 2766 -59 | IR | TYLCV-IL (DQ144621) | TYLCSaV (AY736854) | **R,G**, m,c,3s | $1,2 \times 10^{-19}$ |
| 2400-2723 | C1, C4, IR | TYLCV-IL (DQ144621) | TYLCV-MId (KJ913683) | R,G,B,M,C,S,**3S** | $1,7 \times 10^{-34}$ |

Six samples collected in Pachino on symptomatic plants from breeding material tested negative. As suggested with symptomatic samples from *Ty-1* resistant plants collected in Ouled-Frej (Morocco), the 6 Pachino samples may have been collected from resistant plants in which the virus amount may be too low to be detected. It is also possible that the supposed TYLCD symptoms were caused by some other stresses such as other plant pathogens, or even pesticides. Finally, it cannot be excluded that these samples are infected with TYLCD-associated begomoviruses that are not detectable with the PCR approach used.

The infection profiles determined with the samples collected in Sicily were compared with those determined from tomato samples collected there before 2015 (Belabess et al., 2015; Davino et al., 2008; Davino et al., 2009) (Supplementary material 2). This comparison was possible because similar PCR approaches were used to determine the infection profiles. However, as the PCR techniques used by Davino et al were not designed to distinguish IL and Mld strains (Davino et al., 2008; Davino et al., 2009), the comparison was done after grouping IL and Mld strain-positive samples - distinguished in our study and in that of Belabess et al. - in a unique TYLCV group, and similarly, after grouping IL/Sar and Mld/Sar recombinants in a unique TYLCV/TYLCSaV group. In 2002, shortly after the invasion of TYLCV, only three types of infection profiles were detected (Fig. 4). Subsequently, the number of infection profiles increased with a maximum of 6 infection profiles in 2004 and 2008. Such highly diverse infection profiles were confirmed from samples collected in Sicily between 2006 and 2009, at the rate of 100 samples per year (Davino et al., 2012). However, in spite of high sampling efforts, samples that were positive only for TYLCV/TYLCSaV recombinants (“TYLCV/TYLCSaV”) were not detected before 2009, whereas such samples were detected from 2012.

**Figure 4:**
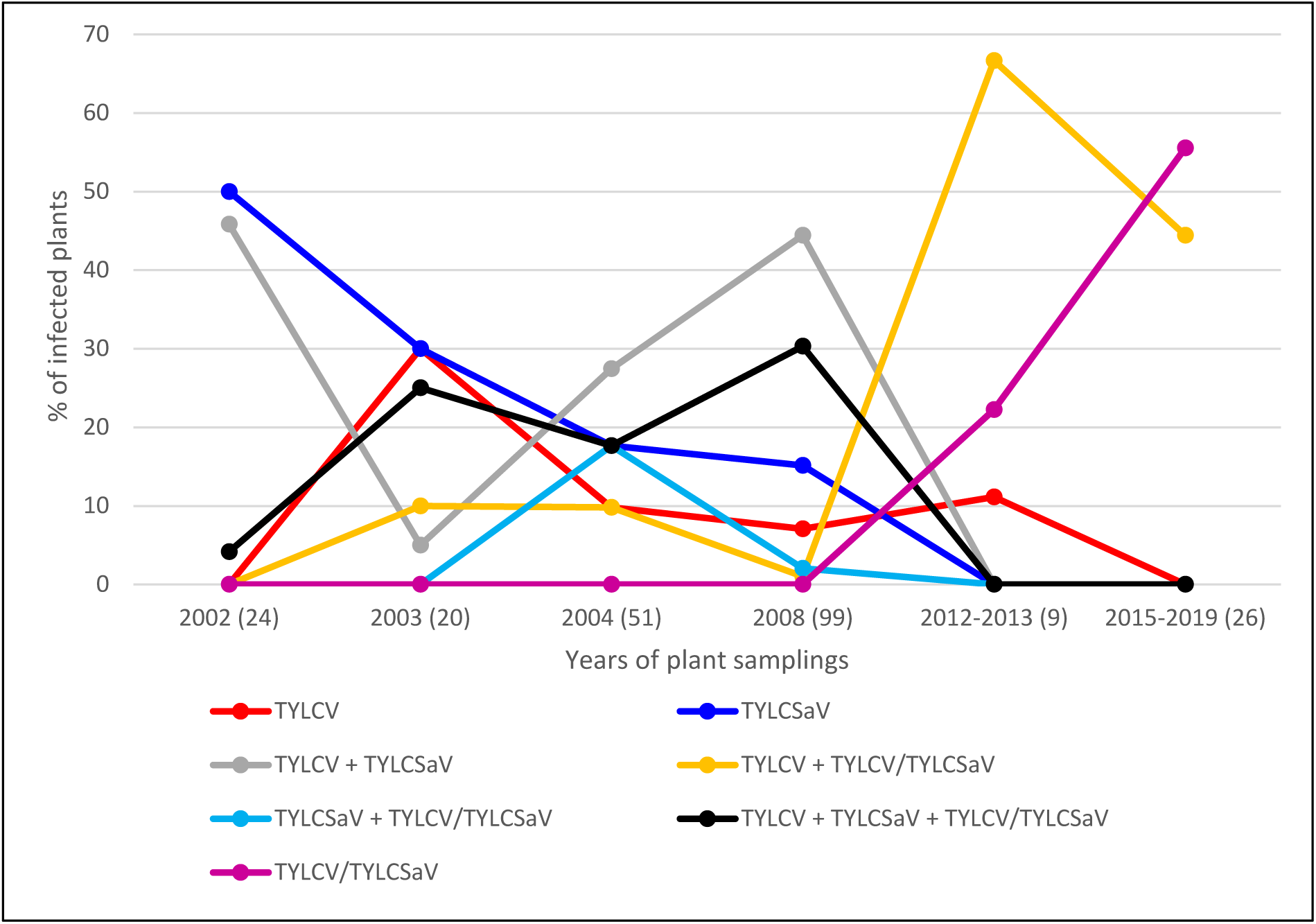
Evolution of the relative frequency of plants exhibiting particular profiles of begomovirus infections in Sicily. The figure presents for each sampling year the percentage of plants corresponding to each infection profile described in the legend. The number of plants analysed in each collection period are indicated in brackets under the graph. The infection profile data of the samples collected between 2015 and 2019 were extracted from Table 1, whereas those of samples collected earlier were extracted from previous reports (Belabess et al., 2015; Davino et al., 2008; Davino et al., 2009). As similar PCR approaches were used in previous and present studies and as all the begomovirus types targeted in the previous studies were also targeted here, it was relevant to present all results on the same graph. The data used to generate Fig. 4 are available in Supplementary material 2. The PCR tool used by Davino et al. was not specific enough to distinguish the two strains of TYLCV (IL and Mld) and their recombinants with TYLCSaV. Hence, for the sake of homogeneity and clarity, the figure does not distinguish the IL and Mld strains for the samples analysed in this study (Table 1) and by (Belabess et al., 2015); IL and Mld strains were combined in a unique TYLCV group and similarly the recombinants IL/Sar and Mld/Sar were combined in a unique TYLCV/TYLCSaV recombinant group. Noteworthy, it is only from the sampling of 2012-2013 that some samples were detected positive only for TYLCV/TYLCSaV recombinants.

### 3.4. Infection profiles of samples collected in Spain

Among the 17 samples analysed, 15 were detected virus-positive and all of these 15 plants exhibited a mono-infection profile. Thus, thirteen samples were positive only for the presence of TYLCV-IL, and two only for TYLCV-IS76-type recombinants. Whereas TYLCV-IL was detected every year of sampling between 2015 and 2018 (13 samples), TYLCV-IS76-type recombinants were detected in 2018 only (2 samples).

The TYLCV-IL-positive samples were analysed further by sequencing a PCR product containing nucleotide positions that can be used to discriminate two TYLCV strains previously reported from Southeast Spain as group 1 and 2 (Torre et al., 2018). Viruses infecting the 5 TYLCV-IL positive plants sampled in 2015 and 2016 clustered with group 1 viruses (Fig. 5); two TYLCV-IL-positive samples collected later (2018) were found to be infected with group 1 viruses too. Viruses infecting all the other TYLCV-IL-positive samples, i.e., 2 collected in 2017 and 3 collected in 2018, clustered with group 2 viruses. Interestingly, one of the three samples collected in 2017 was found to be co-infected with group 1 and group 2 viruses. Indeed, the Sanger-sequencing electropherogram obtained with this sample exhibited double peaks at group1/group2 discriminating positions.

**Figure 5:**
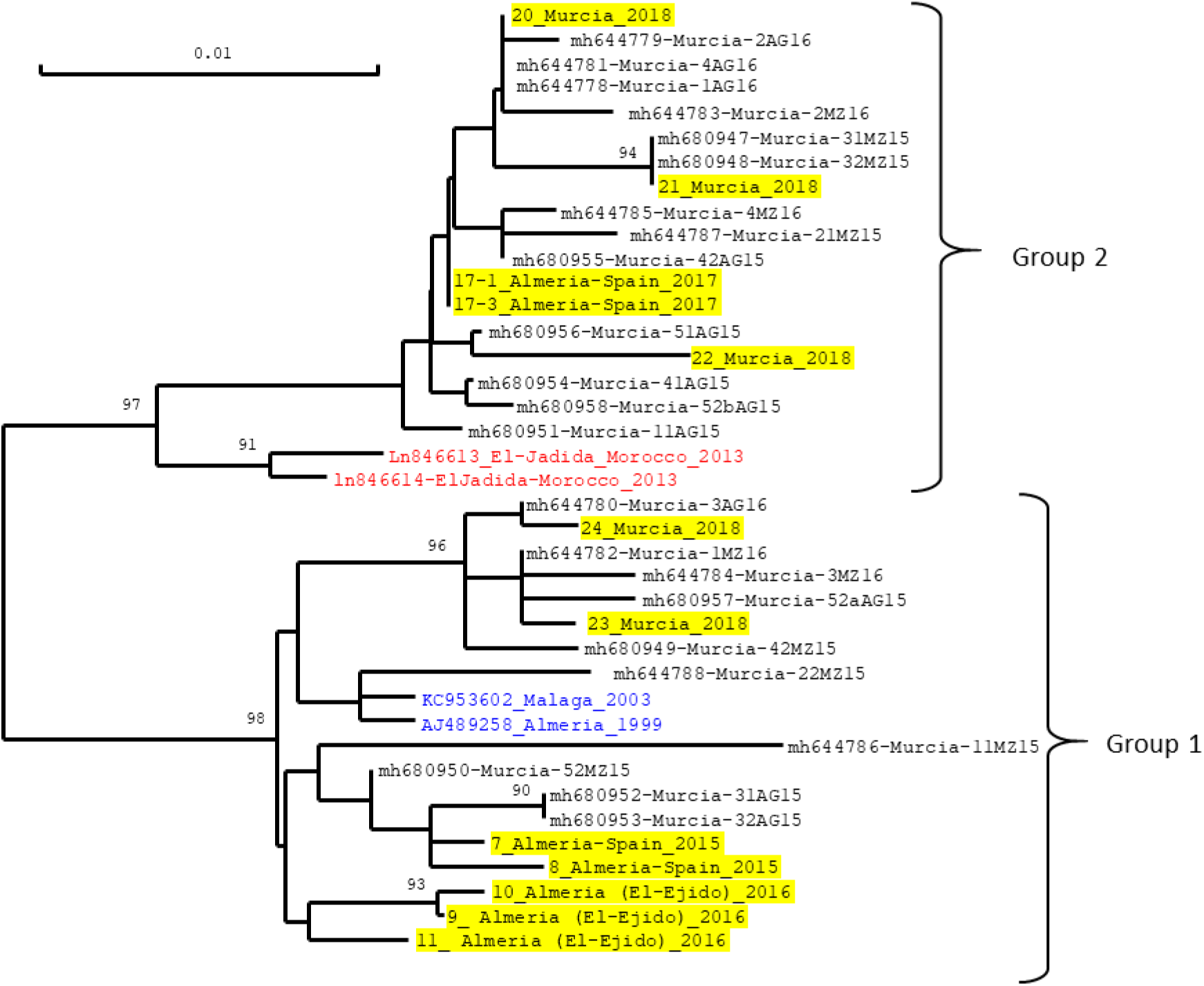
Neighbour-joining tree showing the genetic distance among isolates of Tomato yellow leaf curl virus (TYLCV) originating from Southeast Spain. The tree was constructed from the alignment of a 592-nt-genomic fragment between the origin of replication and the first third of the capsid protein gene. The sequences used for the comparison were from the TYLCV-IL viruses isolated in this study (highlighted in yellow) as well as from other TYLCV-IL isolates for which sequences were downloaded from Genbank. All the isolates were from tomato plants (*Solanum lycopersicum*) except the isolates collected in 1999 and 2003 which were collected from pepper (*Capsicum annuum*) (written in blue). Sequence data from this study were obtained by sequencing a PCR product generated with the primer pair IL-2629F/ IM650R, and the sequences of the 592-nt fragment are available in Supplementary material 3. Virus isolates from this study are identified with a code number followed by the location and year of collection. Isolates written in black without yellow highlights were collected in the province of Murcia and are identified with their GenBank accession number and the code numbers used previously, in which 15 and 16 correspond to the year of collection (2015, 2016) and AG and MZ to the sampling locations, i.e., Aguilas and Mazaron, respectively (Torre et al., 2018). Isolates written in blue correspond to the earliest collected TYLCV in Southeast Spain for which total genome sequences were reported. Isolates written in red correspond to the two TYLCV isolates from El-Jadida (Morocco) that were reported to be the most closely related to group 2 isolates identified as a new type of TYLCV isolates in Southeast Spain (Torre et al., 2018). The scale measures the Jukes and Cantor distance between sequences. Numbers associated with nodes represent the percentage of 1000 bootstrap iterations supporting the nodes; only percentages ≥ 90% are represented. The two highly supported groups correspond to the previously identified group 1 and group 2 (Torre et al., 2018).

Like the 4 samples collected in Ouled-Frej (Morocco) that tested negative in spite of their collection on symptomatic plants, 2 samples collected in Murcia in Oct 2018 revealed the same inconsistency (Table 1). As suggested for the Ouled-Frej samples, the sensitivity of the PCR tests may have been insufficient to detect the potentially lower virus amount that is expected in resistant plants. It is also possible that the milder TYLCD symptoms exhibited in *Ty-1* resistant plants have been confused with symptoms caused by other plant pathogens or pesticides. Finally, it cannot be excluded that these samples are infected with TYLCD-associated begomoviruses that are not detectable with the PCR approach used.

## 4. Discussion

Begomovirus populations associated with TYLCD in western Mediterranean countries are composed of viruses belonging to the alien species *Tomato yellow leaf curl virus*, the native species *Tomato yellow leaf curl Sardinia virus*, and various interspecies recombinants. Although representatives of each of these three TYLCD-associated virus types were reported from Spain, Italy and Morocco, country specificities were previously detected with plant samples collected over time but mostly before 2015; sampling efforts extended up to 2003 in Spain (Garcia-Andres et al., 2007), up to 2008 in Italy (Davino et al., 2009) and up to 2014 in Morocco (Belabess et al., 2015). Considering geographic proximity and trade activities between these countries, as well as the efficient whitefly transmission of these viruses, it was not known if these specificities would be maintained. This question was addressed here by analysing the infection profile of samples collected between 2015 and 2019 in the three countries.

Due to their resistance breaking feature in Ty1 resistant plants (Belabess et al., 2015; Fortes et al., 2023; Panno et al., 2018), Srec recombinants were given special attention by distinguishing them with an original PCR test from Lrec recombinants commonly reported from Spain and Italy (Davino et al., 2012; Garcia-Andres et al., 2006; Monci et al., 2002). Thus, together with the previously reported multiplex PCR tests (Belabess et al., 2015), we were able to specifically determine the proportion of samples containing Srec recombinants among the samples detected positive for TYLCD-associated begomoviruses.

The symptomatic samples that tested negative with all the PCR tests used reveal the potential limitation of the a priori PCR approach. Indeed, besides the possibility of insufficient sensitivity of the PCR tests and confusing symptoms, it cannot be excluded that potential TYLCD-associated viruses were not detectable with the primer sets used. Higher sequencing investments are needed to detect potential TYLCD-associated viruses that may have been missed by the PCR approach used. Such missing risk is even higher with recombinant genomes as shown with TYLCV-IMS60-2400. Indeed, from the sequence of its complete genome, its amplification with MxPCR1 primers is a posteriori quite puzzling because it has a TYLCV-IL-derived sequence at the nucleotide positions targeted by the TYLCV-Mld-specific primer, and a TYLCV-Mld-derived sequence at the nucleotide positions targeted by the TYLCV-IL specific primers.

### 4.1. TYLCV-IS76 is virtually the only TYLCD-associated begomovirus on the Atlantic coast of Morocco

The dominance of Srec recombinants in the samples collected from Morocco and their identification as TYLCV-IS76 is fully consistent with previous results (Belabess et al., 2015). Indeed, these authors showed that TYLCV-IS76 has replaced its parental viruses in the region of Agadir (Souss) and that it started to spread northwards. The northward spread is confirmed here with Srec recombinants being the only begomovirus type detected in the 10 farms located in the north of Agadir along the Atlantic coast up to Casablanca. The detection of TYLCSaV in 2015 in Agadir is consistent with previous results showing that TYLCSaV was more frequently detected than TYLCV among the rare tomato samples in which parental type viruses were detected after the invasion of TYLCV-IS76 (Belabess et al., 2015). Interestingly the “original” group 2-type virus from Morocco (Belabess et al., 2015) that was detected from two tomato samples collected in 2013 in the area surveyed in this study (El Jadida on the Atlantic Coast) was not detected again from the Moroccan samples, whereas such viruses were detected from Spain (Table 2). This striking difference of group 2 virus prevalence between the two countries suggest that the dominance of TYLCV-IS76 has blocked its spread in Morocco. Therefore, considering the relative competitiveness determined experimentally between TYLCV-IS76 and group 2 viruses (Torre et al., 2018), it seems that the relatively higher fitness of group 2 viruses in single infection of *Ty-1*-resistant plants could not compensate the relatively higher fitness of TYLCV-IS76 in coinfection in susceptible and *Ty-1* resistant plants. Finally, further studies are needed with plant samples from northern and eastern Morocco to complete the picture of tomato begomoviruses in Morocco.

### 4.2. Major shift in Sicilian populations of TYLCD-associated begomoviruses between 2009 and 2012

The comparison of the infection profiles of tomato plants sampled over time in Italy (Fig. 4) reveals two important changes that occurred between 2009 and 2012. Firstly, whereas TYLCV/TYLCSaV recombinants were never found in any plant without parental viruses before 2010 in spite of important sampling efforts with 100 plants per year from 2006 to 2009 (Davino et al., 2012), recombinants were found alone in plants sampled after 2011. Indeed among the 9 samples collected in the 2012-2013 period, 2 samples were positive only for TYLCV/TYLCSaV recombinants (Belabess et al., 2015) and among samples collected later (2015-2019) and analysed in this study, “TYLCV/TYLCSaV” is the dominant infection profile (15 samples out of 27, Supplementary material 2). This evolution of infection profiles over time suggests that recombinants with a relatively higher fitness have emerged. The second change that occurred between 2009 and 2012 is the emergence of Srec recombinants. Indeed, while the recombinants reported from Sicily before 2012 were all of the Lrec type (Davino et al., 2012; Davino et al., 2009), the recombinants detected in the samples collected during 2015-2019, were all of the Srec type (Table 2) and 5 out of the 9 samples collected in 2012-2013, were positive for Srec too (Belabess et al., 2015). The two changes observed since 2012, i.e. the occurrence of plant detected positive only for TYLCV/TYLCSaV recombinants and the emergence of Srec recombinants, are presumably linked. Indeed, the relatively higher fitness experimentally assessed for Srec recombinants (Belabess et al., 2016; Jammes et al., 2023) and particularly TYLCV-IS141 (Urbino et al., 2020) - the Srec recombinant frequently detected in Sicily (Belabess et al., 2015; Granier et al., 2019; Panno et al., 2018) - is consistent with its independent spread. As a reminder, the higher fitness of TYLCV-IS141 was deduced from its relatively higher accumulation in susceptible and *Ty-1* resistant plants as comparted to TYLCV-IL, the most competitive parental virus in *Ty-1-*resistant plants (Urbino et al., 2020).

### 4.3. TYLCV-IMS60-2400, a new Srec recombinant spreading in Sicily

Interestingly, besides TYLCV-IS141, a second Srec recombinant was detected in Sicily during the period 2015-2019, i.e., TYLCV-IMS60-2400. Although TYLCV-IMS60-2400 was detected less frequently than TYLCV-IS141 (Table 1), it exhibits features of an emerging virus. Indeed, the 5 samples that were positive for recombinants bearing the RB60 recombination breakpoint were not positive for any of the targeted begomoviruses, indicating that these recombinants are already circulating autonomously. Moreover TYLCV-IS60-type recombinants were detected from samples collected in three of the five years of sampling, and it was detected in at least two distinct collection sites, namely Marina di Ragusa and Vittoria.

### 4.4. Emergence of TYLCV-IS76 and its presumed back recombinant in Southeast Spain

The dominant infection profile of the samples collected in the Murcia and Almeria provinces is “TYLCV-IL” with 13 samples being positive for TYLCV-IL only, among the 15 PCR-positive samples (Table 2). This result is consistent with previous reports with samples collected earlier from the same Spanish region. Thus, the five tomato samples collected in 2013 in Almeria province and analysed with the same PCR approach were PCR positive only for TYLCV-IL (Belabess et al., 2015). Likewise, among 370 tomato samples collected in the two Spanish provinces in 2003 and detected positive for tomato begomoviruses by tissue blot hybridization, 368 were positive only with the DNA probe specific to TYLCV; 2 were positive with both the TYLCV and the TYLCSaV probes and no sample was detected positive for TYLCSaV only (Garcia-Andres et al., 2007). In this later study in which tomato samples were also analysed from the Malaga province, 45 tomato samples from the three provinces, detected positive by tissue-blot hybridization, were selected to be analysed further by sequencing a PCR product comprising the non-coding intergenic region and the 5′-proximal parts of Rep and V2 open reading frames. Among them, only three samples bore TYLCV/TYLCSaV recombinant genomes, two which inherited the TYLCV fragment from a Mld strain parent and one from an IL strain parent. This low proportion of recombinants stands in contrast with the 2 samples detected positive for IL/Sar recombinants among the 15 PCR positive samples analysed here. The much higher prevalence of TYLCV/TYLCSaV recombinants in tomato is apparently related with the length of the TYLCSaV fragment inherited by the recombinants. Whereas the 2 recombinants detected in 2018 have inherited a short 76 nts fragment of TYLCSaV, like that of the previously reported TYLCV-IS76 viruses, the three recombinants detected in 2003 inherited a much longer TYLCSaV fragment that extended beyond the initiation codon of the V2 gene. While the TYLCV-IS76-positive sample form El Ejido confirms the emergence of TYLCV-IS76 in the Almeria province (Fortes et al., 2023; Torre et al., 2019), the detection of TYLCV-IS76 in a sample from Murcia shows that its distribution is not limited to the Almeria province (Table 2).

The high nucleotide identity between TYLCV-IS76 recombinants from Spain and Morocco (Fortes et al., 2023; Torre et al., 2019) suggests that their presence in Spain is due to an introduction from Morocco where it was initially detected in tomato plants sampled in 2010. Such a scenario may also explain the presence of a new type of TYLCV-IL viruses in Southeast Spain because of their close nucleotide identity to TYLCV-IL viruses detected in Morocco from tomato plants sampled in 2013 in El Jadida (Fig. 5). Based on a high sequence identity with the TYLCV-inherited region of TYLCV-IS76, El Jadida-TYLCV-IL-like viruses were presumed to derive from a TYLCV-IS76 recombinant through a back recombination event (Belabess et al., 2015; Torre et al., 2018). Their detection in more than 50% of the TYLCV-IL-positive samples collected in 2017 and 2018, confirm their emergence in Southeast Spain since their first detection from tomato samples collected in 2015 and 2016 (Torre et al., 2018). As these authors have analysed only samples from the province of Murcia, it was not known if the El Jadida-TYLCV-IL-like viruses have emerged elsewhere. Here, with samples collected in 2017 in the province of Almeria (Table 2), we show that their presence is not limited to the province of Murcia. The successful emergence of viruses of the El Jadida-TYLCV-IL-type is consistent with their fitness advantage shown with a Spanish clone inoculated in *Ty-1* resistant tomato plants (Torre et al., 2018). Although this fitness advantage was only observed in single infection but not in co-infection where TYLCV-IS76 is the most competitive in both susceptible and *Ty-1* resistant plants, the emergence of El Jadida-TYLCV-IL-type viruses was possible in Spain presumably because of the relatively low prevalence of TYLCV-IS76. This assumption is consistent with the failure of its emergence in the region of El-Jadida where it was initially reported but where TYLCV-IS76 is the dominant TYLCD-associated begomovirus.

### 4.5 Concluding remarks

The populations of TYLCD-associated begomoviruses that are described in this study reveal that geographic specificities were still present between 2015 and 2019. The specificities concerning parental type viruses (TYLCV, TYLCSaV) is their virtual absence in the sampled region of Morocco along the Atlantic coast, the contrasted situation in Italy where TYLCSaV was detected in Sardinian samples but not in Sicilian samples, and frequent detection of TYLCV in Spain. The specificities concerning recombinants is revealed by the dominance of TYLCV-IS76 in Morocco but no detection of TYLCV-IS141, the dominance of TYLCV-IS141 in Italy but no detection of TYLCV-IS76, and by the confirmation of the emergence of TYLCV-IS76 and the El Jadida-TYLCV-IL-type in Southeast Spain.

However, besides these country specificities, a general trend was revealed with the original PCR test developed here to distinguish Srec and Lrec recombinants. Indeed, while TYLCV/TYLCSaV recombinants detected in tomato samples collected before 2010 were all of the Lrec type (Belabess et al., 2015; Davino et al., 2008; Davino et al., 2012; Davino et al., 2009; Garcia-Andres et al., 2007; Garcia-Andres et al., 2006; Monci et al., 2002), all the tomato samples detected positive for TYLCV/TYLCSaV recombinants in this study are positive for Srec type recombinants and negative for Lrec. Thus, since the first detection of Srec type from Morocco in 2010 and from Sicily in 2012 (Belabess et al., 2015), the prevalence of Srec recombinants has increased at the expenses of Lrec recombinants.

The higher competitiveness of TYLCV-IS76 as compared to Lrec recombinants observed in controlled conditions using vector transmission (Jammes et al., 2023) is consistent with these inverse dynamics of prevalence of Srec versus Lrec recombinants. The large regional scale at which this trend is detected may be explained by the cultivar switch that occurred in the three countries from susceptible to *Ty-1* resistant cultivars, as reported from Morocco (Belabess et al., 2015; Belabess et al., 2016). More samplings and originating from larger areas of the surveyed countries, are needed to confirm and refine the population shifts detected here. More generally, with such a new survey, the evolution of TYLCD-associated begomoviruses may be monitored further in these countries and particularly the dynamics of TYLCV-IS76 and El Jadida-TYLCV-IL-type viruses in Spain, the fate of the new Srec recombinant circulating in Sicily (TYLCV-IMS60-2400), and the potential persistence of parental type viruses in the less intensive tomato production areas such as the North East of Morocco. Finally, to gain a more comprehensive picture of TYLCD-associated virus populations, metagenomic approaches may be used to address the risk of missing viruses that may not be detectable with the PCR approaches as suggested with some symptomatic plants sampled in this study and the relatively complex recombination profile of TYLCV-IMS60-2400.

## Acknowledgments

We thank Mme Amal Smaili for her assistance in the collection of Moroccan samples. We thank Sylvie Dallot for her assistance in the use of RDP and SimPlot applications.

## Author contributions

Mohamed Faize (MF) organized the collection of Moroccan samples. Martine Granier (MG) and Sandie Passera extracted total DNA from plant samples. MG performed the laboratory analysis of the samples. Michel Peterschmitt (MP) designed the study, analysed the data and wrote the original draft. MF, Cica Urbino and MP reviewed and edited the manuscript.

## Data

DNA sequences of the three complete genomes, generated in this study for the newly reported TYLCV-IMS60-2400 recombinant, are available in GenBank under accession numbers OR473055, OR473056, OR473057. The partial sequences of other viruses generated in this study and used to construct the distance tree shown in Fig. 5 are available in the Supplementary material 3. The data used to prepare Fig. 4 are available in Supplementary material 2.

## Supplementary material 1

Geographical distribution of begomoviruses associated with tomato yellow leaf curl disease, detected in samples collected from Spain Italy and Morocco.

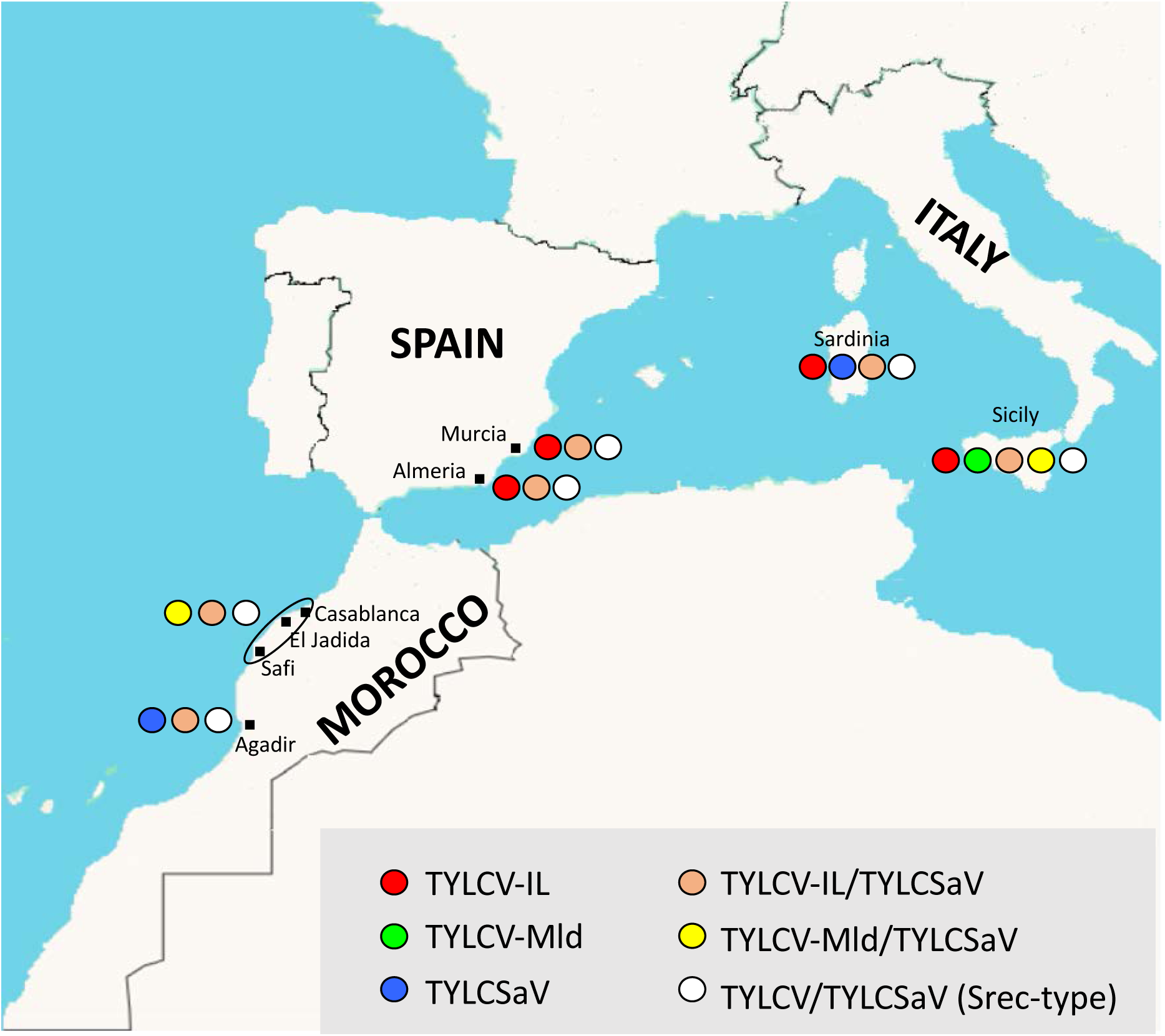

## Supplementary material 2

The data presented in this table were used to generate Fig. 4. They present the number of tomato plants of each infection profiles sampled in different years in Sicily. The data obtained with the plants sampled in the years 2002 and 2004 are from [(Davino et al., 2008), Table 2], those from years 2004 and 2008 are from[(Davino et al., 2009), Table 2], those from years 2012-2013 are from [(Belabess et al., 2015), Supplementary Table S8], and those from years 2015-2019 are from this study (Table 1). Table cells corresponding to plants detected positive for TYLCV/TYLCSAV recombinants are highlighted in orange. The penultimate table line shows the total number of plants detected positive regardless of their infection profile.

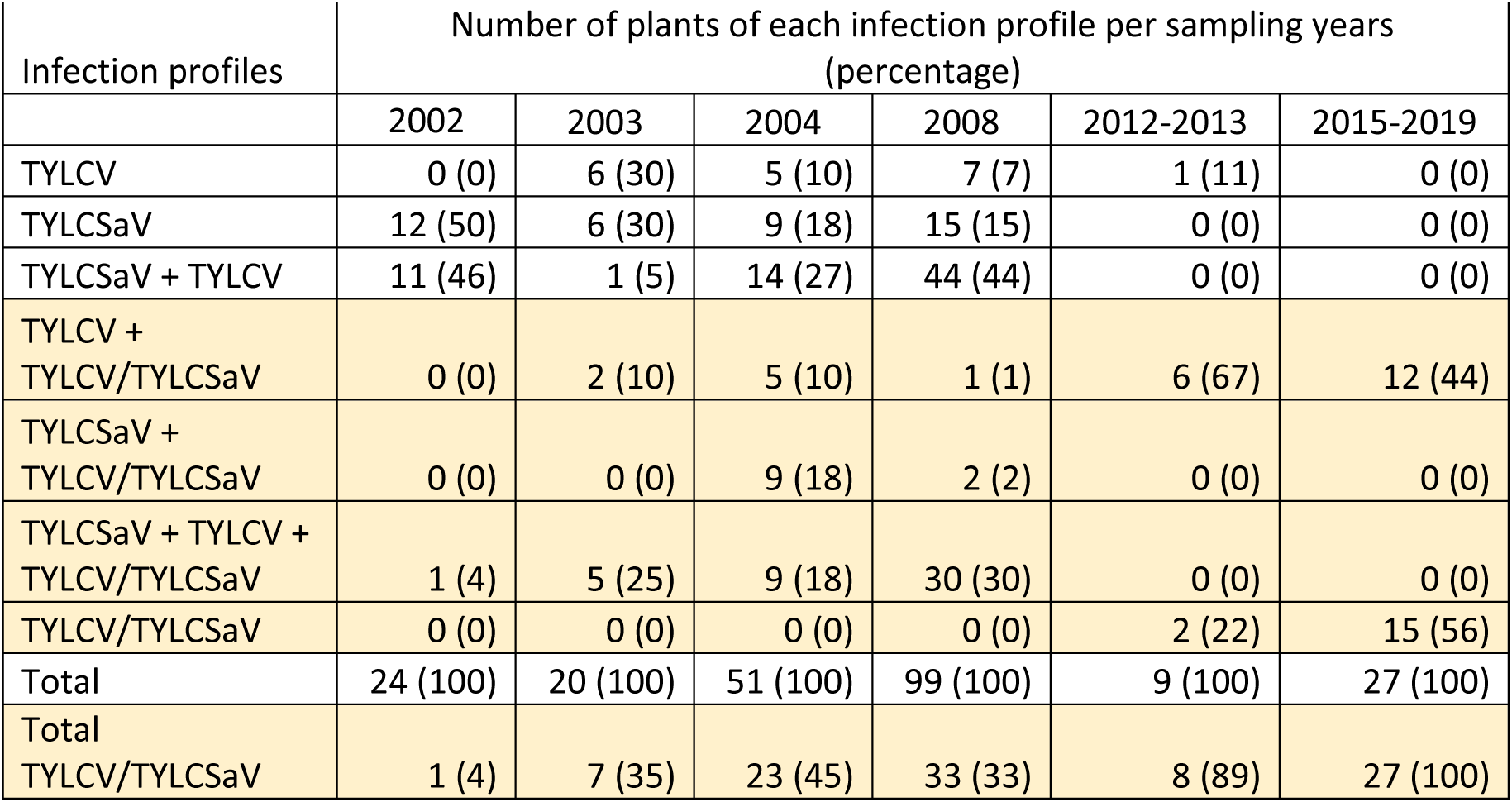

## Supplementary material 3

DNA sequences of the 592-nt-fragment with which the 13 TYLCV-IL isolates detected in tomato samples collected in Southeast Spain (Table 2) were identified as belonging to the previously reported group 1 or group 2 (Torre et al., 2018) (Fig. 5). Only 12 of them are presented, because one of the three samples collected in the province of Almeria in 2017 (Table 2) was co-infected with group 1 and group 2 viruses; indeed, the electropherogram revealed double peaks at nucleotide positions at which the two types of viruses can be distinguished.

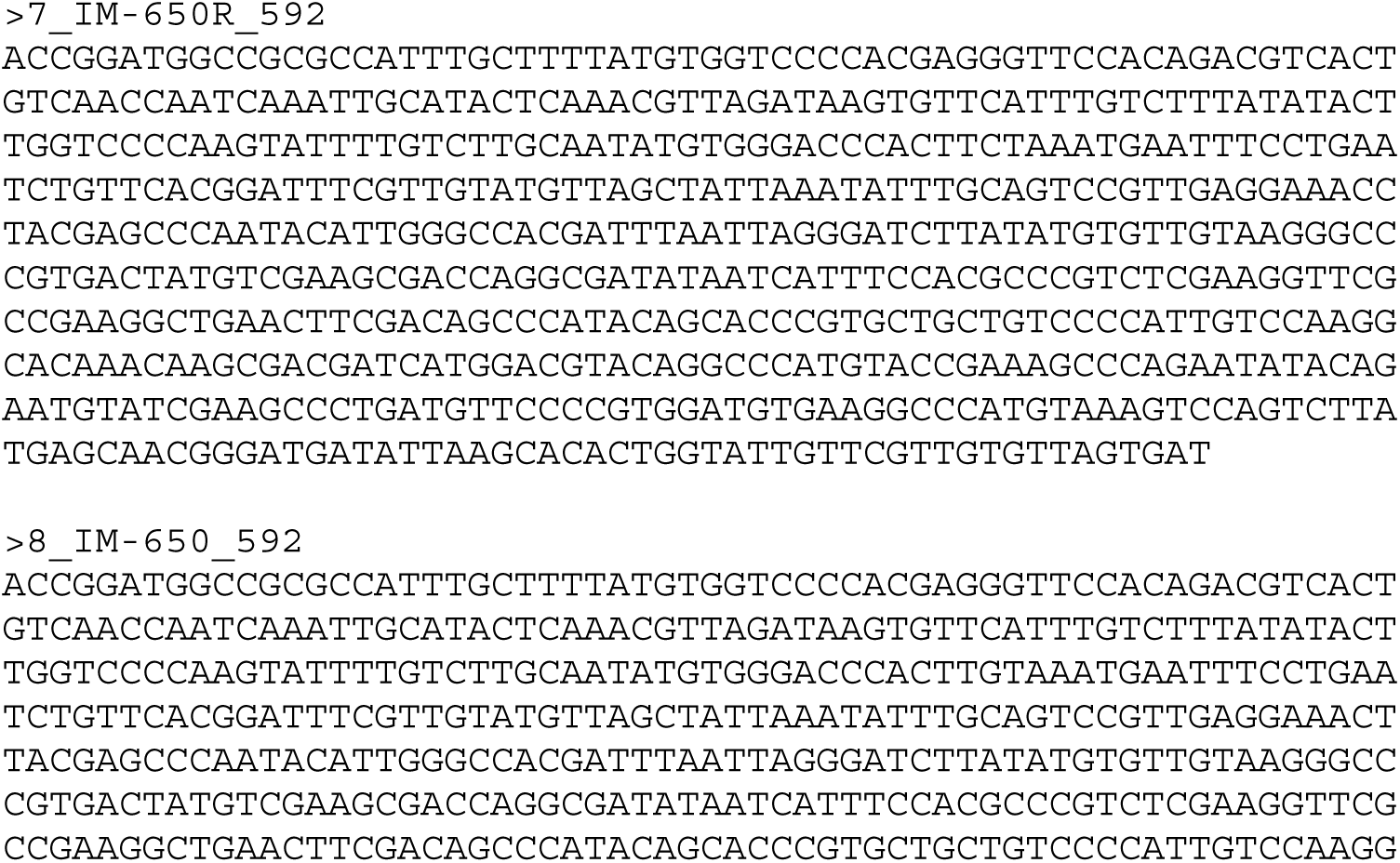
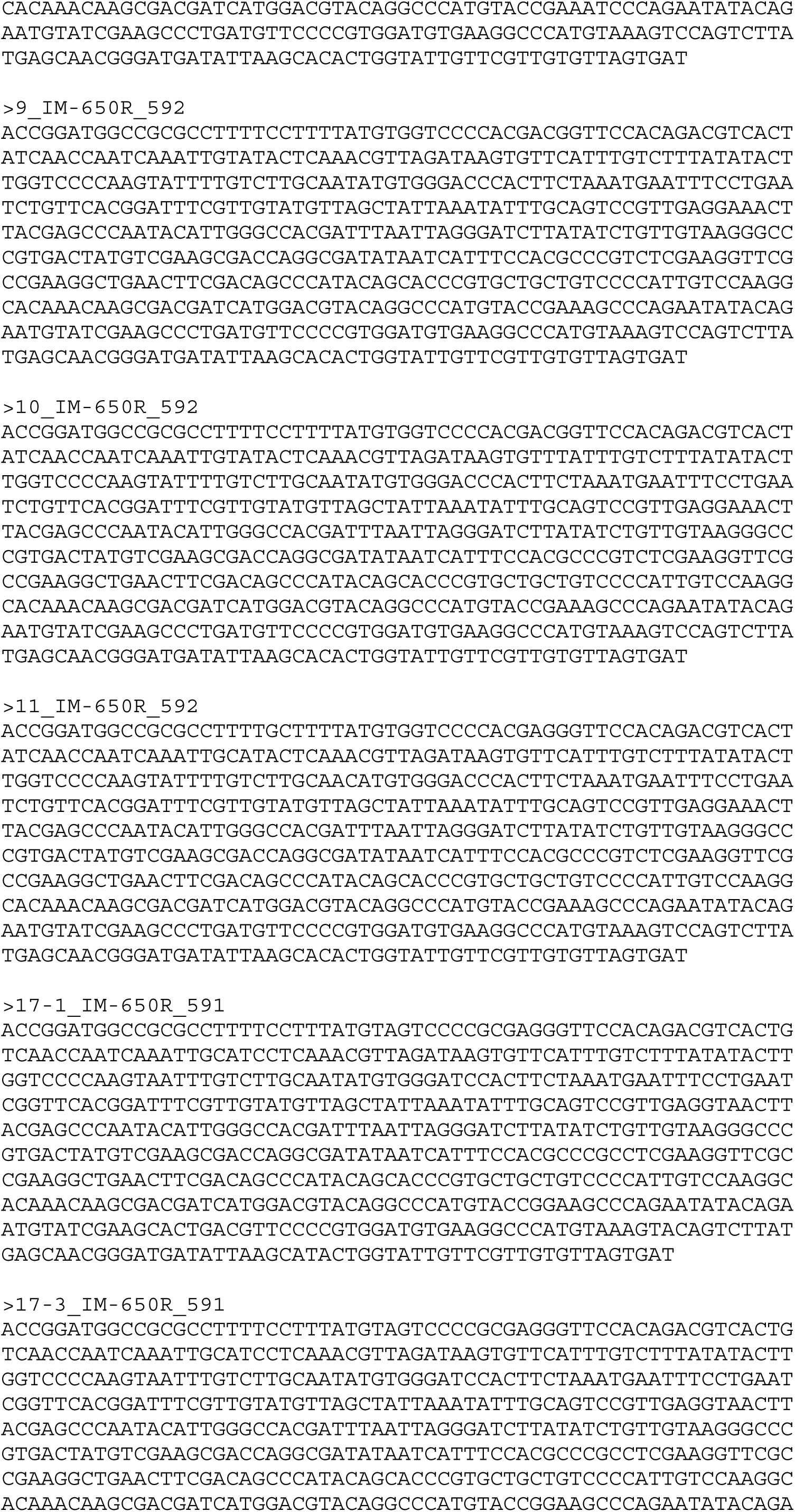
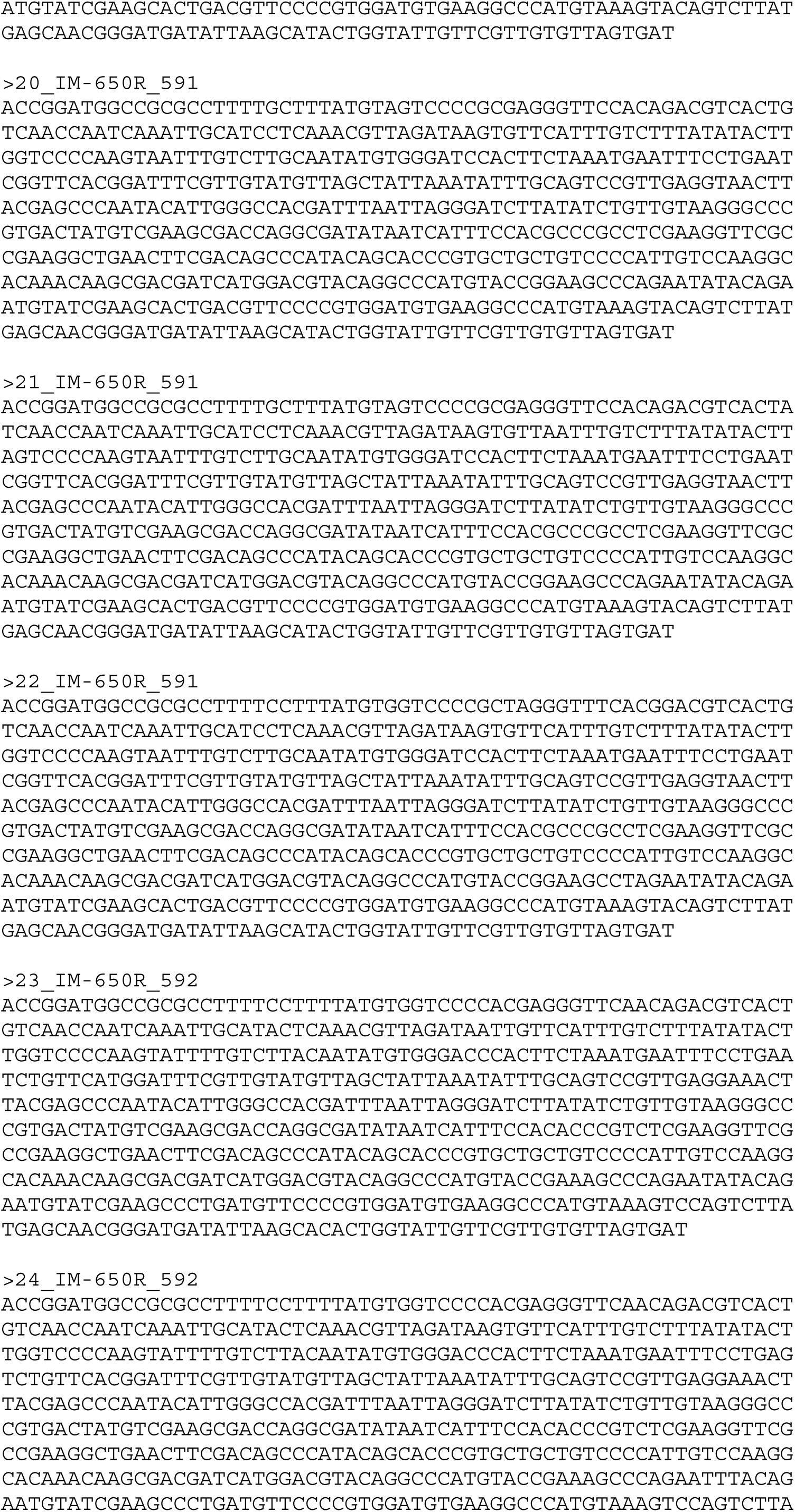
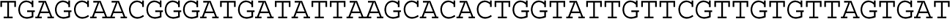

